# Benchmark of tools for CNV detection from NGS panel data in a genetic diagnostics context

**DOI:** 10.1101/850958

**Authors:** José Marcos Moreno-Cabrera, Jesús del Valle, Elisabeth Castellanos, Lidia Feliubadaló, Marta Pineda, Joan Brunet, Eduard Serra, Gabriel Capellà, Conxi Lázaro, Bernat Gel

## Abstract

**Motivation:** Although germline copy number variants (CNVs) are the genetic cause of multiple hereditary diseases, detecting them from targeted next-generation sequencing data (NGS) remains a challenge. Existing tools perform well for large CNVs but struggle with single and multi-exon alterations. The aim of this work is to evaluate CNV calling tools working on gene panel NGS data with CNVs up to single-exon resolution and their suitability as a screening step before orthogonal confirmation in genetic diagnostics strategies.

**Results:** Five tools (DECoN, CoNVaDING, panelcn.MOPS, ExomeDepth and CODEX2) were tested against four genetic diagnostics datasets (495 samples, 231 CNVs), using the default and sensitivity-optimized parameters. Most tools were highly sensitive and specific, but the performance was dataset-dependant. In our in-house datasets, DECoN and panelcn.MOPS with optimized parameters showed enough sensitivity to be used as screening methods in genetic diagnostics.

**Availability:** Benchmarking-optimization code is freely available at https://github.com/TranslationalBioinformaticsIGTP/CNVbenchmarkeR.

## Introduction

Next-generation sequencing (NGS) is an outstanding technology to detect point mutations and small deletion and insertion variants in genetic testing for Mendelian conditions. However, detection of large rearrangements such as copy-number variants (CNV) from NGS data is still challenging due to issues intrinsic to the technology including short read lengths and GC-content bias (Teo *et al*., 2012). Nevertheless, it is well recognized that germline CNVs are the genetic cause of several hereditary diseases (Zhang *et al*., 2009), so their analysis is a necessary step in a comprehensive genetic diagnostics strategy.

The gold standards for CNV detection in genetic diagnostics are multiplex ligation-dependent probe amplification (MLPA) and array comparative genomic hybridization (aCGH) (Kerkhof *et al*., 2017). Both methods are time-consuming and costly, so frequently only a subset of genes is tested, excluding others from the analysis, especially when using single-gene approaches. Therefore the possibility of using NGS data as a first CNV screening step would decrease the number of MLPA/aCGH tests required and would free up resources.

Many tools for CNVs detection from NGS data have been developed (Zhao *et al*., 2013; Abel and Duncavage, 2013; Mason-Suares *et al*., 2016). Most of them can reliably call large CNVs (in the order of megabases) but show poor performance when dealing with small CNVs affecting only one or few small exons, which are CNVs frequently involved in several genetic diseases (Truty *et al*., 2019). In addition, most of these tools were designed to work with whole genome or whole exome data and struggle with the sparser data from NGS gene panels used in routine genetic testing. Therefore, the challenge is to identify a tool able to detect CNVs from NGS panel data at a single exon resolution with sufficient sensitivity to be used as a screening step in a diagnostic setting.

Other benchmarks of CNV calling tools on targeted NGS panel data have been published. However, they were performed by the authors of the tools and executed against a single dataset(Johansson *et al*., 2016; Fowler *et al*., 2016; Povysil *et al*., 2017; Kim *et al*., 2017; Chiang *et al*., 2019), or used mainly simulated data with a small number of validated CNVs (Roca *et al*., 2019). The aim of this work is to perform an independent benchmark of multiple CNV calling tools, optimizing and evaluating them against multiple datasets generated in diagnostics settings, to identify the most suitable tools to be used for genetic diagnostics (Figure 1).

**Figure 1.**
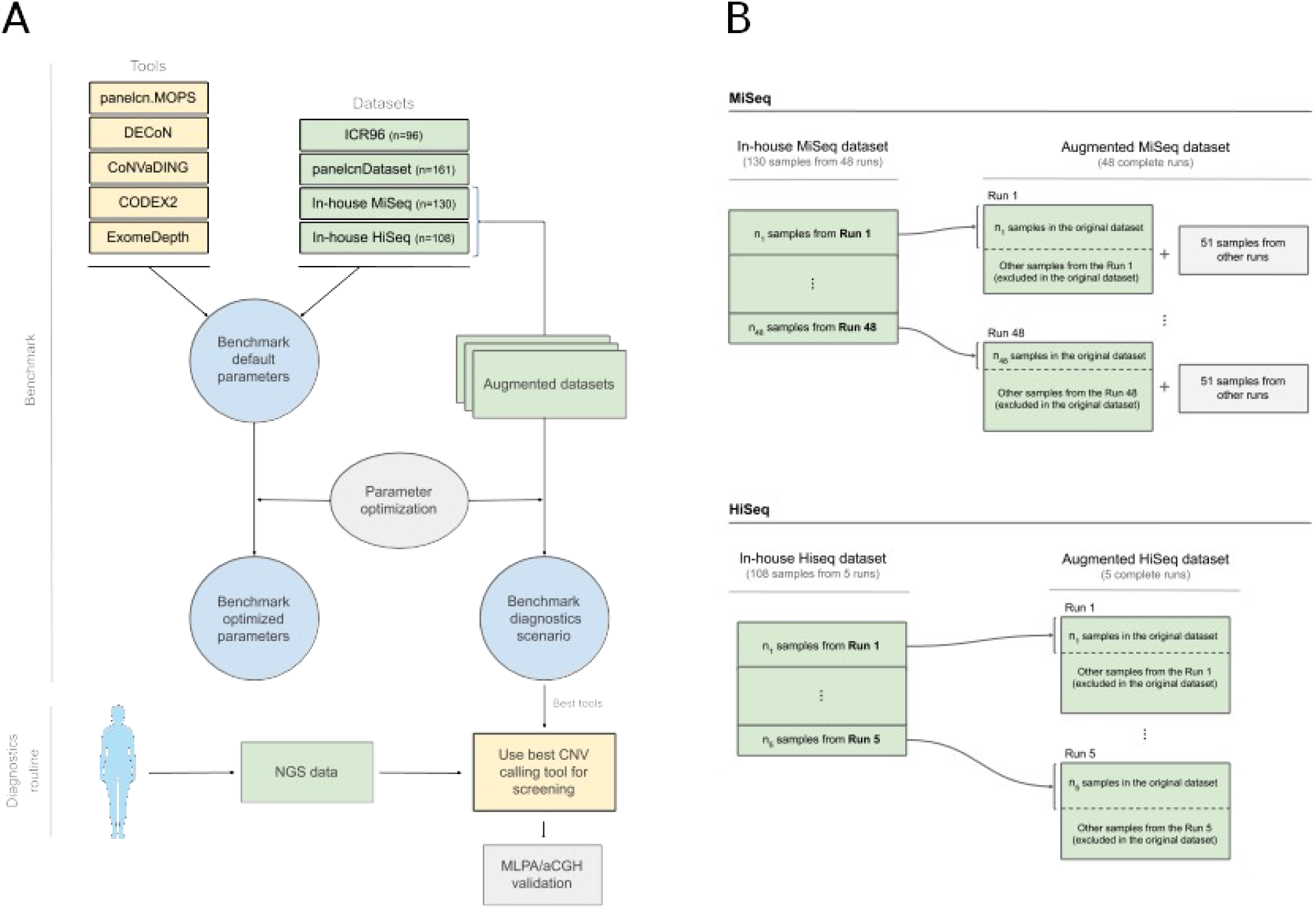
Benchmark design and augmented datasets. (**A**) The panel shows the benchmark design and the objective of applying the results in the diagnostics routine. (**B**) To evaluate the diagnostics scenario, a new dataset was built for each run belonging to the original dataset. The augmented datasets contained all the samples originally sequenced in the run and, in the case of the MiSeq datasets (upper), a set of 51 samples with no known CNV from different runs.

## Methods

### Datasets

Four datasets were included in this benchmark (ICR96, panelcnDataset, In-house Miseq and In-House HiSeq) (Table 1) with data from hybridization-based target capture NGS panels designed for hereditary cancer diagnostics (TruSight Cancer Panel (Illumina) and I2HCP (Castellanos *et al*., 2017). All datasets were generated in real diagnostics settings and contained single and multi-exon CNVs, all of them validated by MLPA. Negative MLPA data, meaning no detection of any CNV, was also available for a subset of genes.

**Table 1:** NGS panel datasets

|  | **Samples** | **Validated genes with CNV** | **Single-exon CNVs** | **Multi-exon CNVs** | **Validated genes with no CNV** | **Sequencing** | **Additional info** |
| --- | --- | --- | --- | --- | --- | --- | --- |
| ICR96 | 96 | 68 | 25 | 43 | 1752 (96.3% of total) | TruSight Cancer Panel v2 (100 genes) HiSeq | Samples obtained from one run |
| panelcnData set | 161 | 41 | 13 | 28 | 416 (91% of total) | TruSight Cancer Panel (94 genes) MiSeq |  |
| In-house MiSeq | 130 | 64 | 19 | 45 | 167 (72.3% of total) | I2HCP Panel (122-135 genes) MiSeq | Samples obtained from 48 runs. Three samples had a CNV in mosaicism |
| In-house HiSeq | 108 | 58 | 18 | 40 | 176 (75.2% of total) | Panel I2HCP (135 genes), HiSeq | Samples obtained from 5 runs. Two samples had CNV in mosaicism |

The dataset ICR96 exon CNV validation series (Mahamdallie *et al*., 2017) was downloaded from the European Genome-phenome Archive (EGA) (EGAD00001003335) and contained 96 samples captured with TruSight Cancer Panel v2 (100 genes) and sequenced in a single HiSeq lane with 101-base paired-end reads. panelcnDataset (Povysil *et al*., 2017) was also downloaded from EGA (EGAS00001002481) and had 170 samples captured using TruSight Cancer Panel (94 genes, Illumina) and sequenced on a MiSeq instrument with 151-base paired-end reads. Only 161 out of the 170 samples were used in this work and 9 were removed because they presented alterations out of the scope of this benchmark: 5 presented CNVs smaller than an exon (IBK9, IBK23, IBK67, IBK153, IBK166) and 4 contained ALUs insertions instead of CNVs (IBK141, IBK142, IBK143, IBK151). IBK141 ALU insertion was not reported in the original publication but identified in this work (Supplementary File 1).

The in-house datasets were generated at the ICO-IGTP Joint Program for Hereditary Cancer and are available at the EGA under the accession number (*submission in progress). Both datasets were captured with our custom I2HCP panel v2.0-v2.2 (122-135 genes) and sequenced in either a MiSeq with 2×300 bp reads (In-house Miseq, 130 samples) or a HiSeq with 2×251 bp reads (In-house Hiseq, 108 samples). A total of 1103 additional samples (505 MiSeq, 598 HiSeq), with no CNVs detected in the subset of genes tested by MLPA, were used to build the augmented datasets used in the diagnostics scenario analysis. Informed consent was obtained for all samples in the in-house datasets.

Detailed information on MLPA-detected CNVs for each dataset can be found in supplementary files 4, 5, 6 and 7.

### Tools

Five tools were tested in the benchmark: CoNVaDING v1.2.0 (Johansson *et al*., 2016), DECoN v1.0.1 (Fowler *et al*., 2016), panelcn.MOPS v1.0.0 (Povysil *et al*., 2017), ExomeDepth v1.1.10 (Plagnol *et al*., 2012) and CODEX2 v1.2.0 (Jiang *et al*., 2018).

### Data preprocessing

All samples were aligned to the GRCh37 human genome assembly using BWA mem v0.7.12 (Li and Durbin, 2009; Li, 2013). SAMtools v0.1.19 (Li *et al*., 2009) was used to sort and index BAM files. No additional processing or filtering was applied to the BAM files.

### Regions of interest

The regions of interest (ROIs) were dependent on the dataset. For TruSight based datasets, ICR96 and panelcnDataset, we used the targets bed file published elsewhere (Fowler *et al*., 2016) with some modifications: the fourth column was removed, the gene was added and file was sorted by chromosome and start position (Supplementary File 2). For in-house datasets, we generated a target bed file containing all coding exons from all protein-coding transcripts of genes in the I2HCP panel v2.1 (Supplementary File 3). This data was retrieved from Ensembl Biomart version 67 (Flicek *et al*., 2012) may2012.archive.ensembl.org). All genes tested by MLPA and used in the benchmark were common to all I2HCP versions (v2.0-2.2).

### Benchmark evaluation metrics

The performance of each tool for CNVs detection was evaluated at two levels: per region of interest (ROI) and per gene.

Per ROI metrics treated all regions of interest as independent entities, assigning each of them a correctness value: true positive (TP) or true negative (TN) if the tool matched the results of MLPA, false negative (FN) if the tool missed a CNV detected by MLPA and false positive (FP) if the tool called a CNV not detected by MLPA. This is the most fine-grained metric.

Per gene metrics consider the fact that most MLPA kits cover a whole gene and so the true CNVs would be detected by MLPA when confirming any CNV call in any ROI of the affected gene. Therefore, per gene metrics assigned a correctness value to each gene taking into account all its exons: TP if one of its ROIs was a TP; FN if MLPA detected a CNV in at least one of its ROIs and none of them were detected by the tool; FP if the tool called a CNV in at least one ROI and none of them were detected by MLPA; TN if neither MLPA or the tool detected a CNV in any of its ROIs.

For each tool against each dataset and evaluation level various performance metrics were computed: sensitivity defined as TP / (TP + FN), specificity defined as TN / (TN + FP), positive predictive value (PPV) defined as TP / (TP + FP), negative predictive value (NPV) defined as TN / (TN + FN), accuracy (ACC) defined as (TP + TN) / (TP + FP + FN + TN), F1 score (F1) defined as 2TP / (2TP + FP + FN), Matthews correlation coefficient (MCC) defined as (TP * TN - FP * FN) / ([(TP + FP) * (FN + TN) * (FP + TN) * (TP + FN)] ^ (1/2)) and Cohen’s kappa coefficient defined as (ACC – EACC) / (1 – EACC) where expected accuracy (EACC) is defined as (TP + FP) * (TP + FN) + (TN + FP) * (TN + FN) / ((TN + FP + TP + FN) ^2).

### Parameter Optimization

Parameters of each tool were optimized against each dataset to maximize sensitivity while limiting specificity loss: each dataset was split into two halves, a training set used to optimize tool parameters and a validation set to evaluate them. The optimization algorithm followed a greedy approach: a local optimization was performed at each step with the aim of obtaining a solution close enough to the global optimum. Further details of the optimization algorithm can be found in Supplementary File 8.

### Benchmarking framework execution

An R framework, CNVbenchmarkeR, was built to perform the benchmark in an automatically and configurable way. Code and documentation are available at https://github.com/TranslationalBioinformaticsIGTP/CNVbenchmarkeR. Each selected tool was first executed against each dataset using default parameters as defined in tool documentation and then using the optimized parameters. Default and optimized parameter values can be found in Supplementary File 9.

Tool outputs were processed with R v3.4.2 and Bioconductor v3.5 (Gentleman *et al*., 2004), plyr (Wickham, 2011), GenomicRanges (Lawrence *et al*., 2013) and biomaRt (Durinck *et al*., 2009). Plots were created with ggplot2 (Wickham, 2016).

### Diagnostics scenario evaluation

The In-house MiSeq and In-house HiSeq datasets were composed of a selection of samples from different sequencing runs. In a real diagnostics scenario, the objective is to analyze a new run with all its sequenced samples. To simulate and evaluate the diagnostics scenario, we built the augmented datasets (Figure 1), which contained all the samples from the sequencing runs instead of a selection of them. For the augmented datasets, the tools were executed against each run and metrics were computed by combining the results of all runs. Since some tools recommend more than 16 samples for optimal performance, we added 51 samples from other runs with no known CNVs when executing the tools on the runs of the augmented MiSeq dataset.

We also defined a new metric, whole diagnostics strategy, to take into account that in a diagnostics setting all regions where the screening tool was not able to produce a result (no-call) should be identified and tested by other methods. Thus, any gene containing at least one positive call or no-call in a ROI was considered as a positive call of the whole gene: TP if the gene contained at least one ROI affected by a CNV; FP if the gene did not contain any ROI affected by a CNV.

## Results

To identify the CNV calling tools that could be used as a screening step in a genetic diagnostics setting, we needed first to select the candidate tools, and then to evaluate their performance with a special emphasis on the sensitivity, both with their default parameters and with dataset-dependent optimized parameters.

### CNV calling tool selection

The first in the benchmark was to identify candidate tools that have shown promising results. After a literature search process, we selected five CNV calling tools to be evaluated (Table 2), all of them based on depth-of-coverage analysis. Three tools have been reported to perform well on NGS panel data at single-exon resolution: CoNVaDING v1.2.0 (Johansson *et al*., 2016), DECoN v1.0.1 (Fowler *et al*., 2016) and panelcn.MOPS v1.0.0 (Povysil *et al*., 2017). ExomeDepth v1.1.10 (Plagnol *et al*., 2012) was included due to its high performance in benchmarks on WES data (de Ligt *et al*., 2013; Sadedin *et al*., 2018) and because the developers reported good performance with panel data (https://github.com/vplagnol/ExomeDepth). CODEX2 v1.2.0 was included due to the high sensitivity shown on WES data (Jiang *et al*., 2018) and the availability of specific scripts for panel data (https://github.com/yuchaojiang/CODEX2).

**Table 2:** Summary of CNV calling tools evaluated in this study

|  | Language | Version | Number of parameters used in the benchmark | Detect low quality regions | Availability |
| --- | --- | --- | --- | --- | --- |
| CODEX2 | R package | 1.2.0 <sup>a</sup> | 10 | No | https://github.com/yuchaojiang/CODEX2 |
| CoNVaDING | Perl program | 1.2.0 | 7 | Yes | https://github.com/molgenis/CoNVaDING |
| DECoN | R program | 1.0.1 | 3 | Yes | https://github.com/RahmanTeam/DECoN |
| ExomeDepth | R package | 1.1.10 | 4 | No | https://github.com/vplagnol/ExomeDepth |
| panelcn.MOPS | R package | 1.0.0 | 13 | Yes | https://github.com/bioinf-jku/panelcn.mops |

<sup>a</sup> CODEX2 script for panel setting (Codex2\_targeted.R) was obtained from version dated at on Sep 12, 2017.

### Benchmark with default parameters

We executed each tool on each dataset with the default parameters and computed evaluation statistics at two levels: per region of interest (ROI) and per gene (see Methods).

Regarding the per ROI metric, most tools showed sensitivity and specificity values over 0.75, with sensitivity in general over 0.9 (Figure 2 and Table 3). However, tool performance varied across datasets. For the ICR96 and panelcnDataset datasets, specificity was always higher than 0.98, while sensitivity remained higher than 0.94 (with the exception of CODEX2). This performance was not achieved when using the in-house datasets, where lower sensitivity and specificity can be observed, and only CoNVaDING obtained sensitivity close to 1 at the expense of a lower specificity.

**Table 3.**
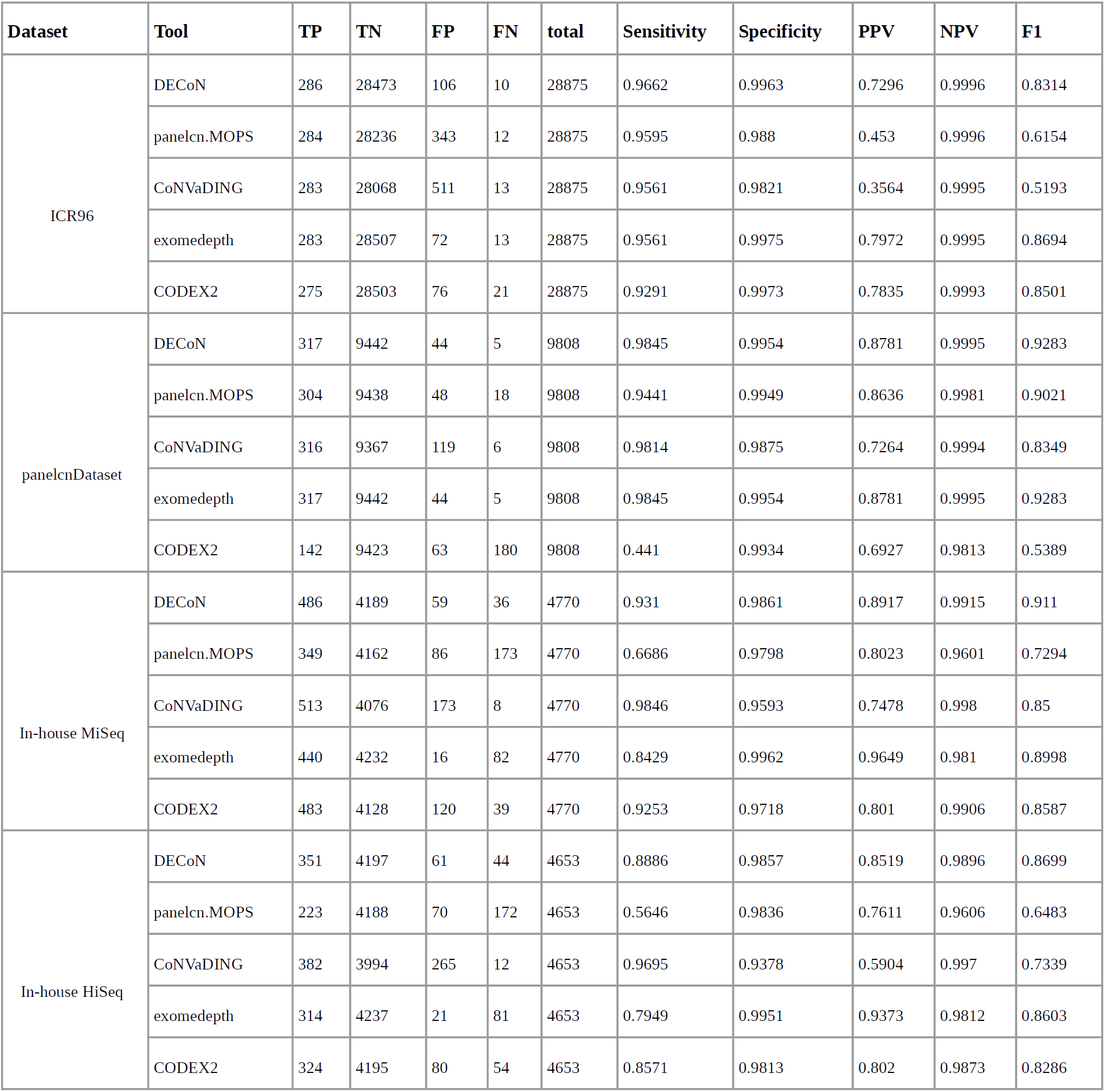
Benchmark results with default parameters: per ROI metrics. Shows results when executing tools with the default parameters and computing the per ROI metrics.

**Figure 2.**
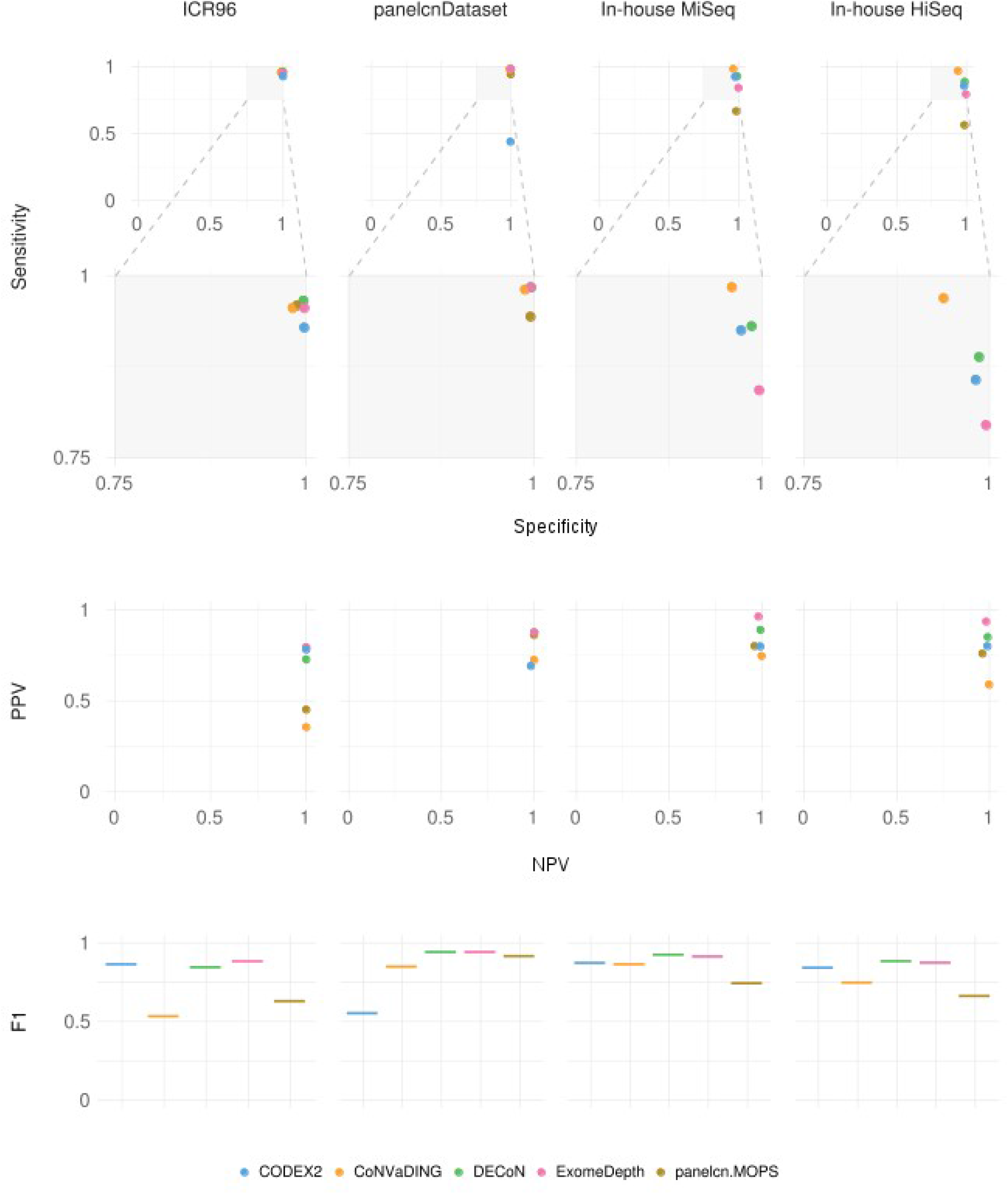
Benchmark results with default parameters: per ROI metrics. Shows results when executing tools with the default parameters and computing the per ROI metrics. ExomeDepth and DECoN tools obtained same sensitivity and specificity in panelcnDataset.

As expected in unbalanced datasets with a much larger number of negative elements than positive ones, negative predictive value (NPV) was higher than positive predictive value (PPV) in all tool-dataset combinations. All NPVs were above 0.96 while PPV varied across datasets, ranging from 0.36 (CoNVaDING in ICR96) to 0.96 (ExomeDepth in In-house MiSeq). ExomeDepth had the highest PPV in all datasets.

Regarding the per gene metric, sensitivity was slightly improved compared to per ROI, and for each dataset, at least one tool showed a sensitivity of 1 and was able to identify all CNVs (Table 4, Supplementary Figure 1 and Supplementary File 10).

**Table 4:** Benchmark results with default parameters: per gene metrics. Shows results when executing tools with the default parameters and computing the per gene metrics.

| Dataset | Tool | TP | TN | FP | FN | total | Sensitivity | Specificity | PPV | NPV | F1 |
| --- | --- | --- | --- | --- | --- | --- | --- | --- | --- | --- | --- |
| ICR96 | DECoN | 63 | 1703 | 49 | 5 | 1820 | 0.9265 | 0.972 | 0.5625 | 0.9971 | 0.7 |
|  | panelcn.MOPS | 68 | 1646 | 106 | 0 | 1820 | 1 | 0.9395 | 0.3908 | 1 | 0.562 |
|  | CoNVaDING | 66 | 1650 | 102 | 2 | 1820 | 0.9706 | 0.9418 | 0.3929 | 0.9988 | 0.5593 |
|  | exomedepth | 62 | 1729 | 23 | 6 | 1820 | 0.9118 | 0.9869 | 0.7294 | 0.9965 | 0.8105 |
|  | CODEX2 | 63 | 1741 | 11 | 5 | 1820 | 0.9265 | 0.9937 | 0.8514 | 0.9971 | 0.8873 |
| panelcnDataset | DECoN | 41 | 407 | 9 | 0 | 457 | 1 | 0.9784 | 0.82 | 1 | 0.9011 |
|  | panelcn.MOPS | 41 | 403 | 13 | 0 | 457 | 1 | 0.9688 | 0.7593 | 1 | 0.8632 |
|  | CoNVaDING | 40 | 405 | 11 | 1 | 457 | 0.9756 | 0.9736 | 0.7843 | 0.9975 | 0.8696 |
|  | exomedepth | 41 | 407 | 9 | 0 | 457 | 1 | 0.9784 | 0.82 | 1 | 0.9011 |
|  | CODEX2 | 37 | 414 | 2 | 4 | 457 | 0.9024 | 0.9952 | 0.9487 | 0.9904 | 0.925 |
| In-house MiSeq | DECoN | 60 | 169 | 3 | 4 | 236 | 0.9375 | 0.9826 | 0.9524 | 0.9769 | 0.9449 |
|  | panelcn.MOPS | 55 | 151 | 21 | 9 | 236 | 0.8594 | 0.8779 | 0.7237 | 0.9438 | 0.7857 |
|  | CoNVaDING | 64 | 152 | 20 | 0 | 236 | 1 | 0.8837 | 0.7619 | 1 | 0.8649 |
|  | exomedepth | 55 | 171 | 1 | 9 | 236 | 0.8594 | 0.9942 | 0.9821 | 0.95 | 0.9167 |
|  | CODEX2 | 59 | 162 | 10 | 5 | 236 | 0.9219 | 0.9419 | 0.8551 | 0.9701 | 0.8872 |
| In-house HiSeq | DECoN | 56 | 175 | 3 | 2 | 236 | 0.9655 | 0.9831 | 0.9492 | 0.9887 | 0.9573 |
|  | panelcn.MOPS | 47 | 158 | 20 | 11 | 236 | 0.8103 | 0.8876 | 0.7015 | 0.9349 | 0.752 |
|  | CoNVaDING | 58 | 161 | 17 | 0 | 236 | 1 | 0.9045 | 0.7733 | 1 | 0.8722 |
|  | exomedepth | 52 | 175 | 3 | 6 | 236 | 0.8966 | 0.9831 | 0.9455 | 0.9669 | 0.9204 |
|  | CODEX2 | 54 | 172 | 6 | 4 | 236 | 0.931 | 0.9663 | 0.9 | 0.9773 | 0.9153 |

### Benchmark with optimized parameters

In addition to evaluating the performance of the different tools tested with default parameters, we performed an optimization process to identify, for each tool and dataset, the combination of parameters that maximized the sensitivity as required for a screening tool in a diagnostics context (see Methods and Supplementary <u>File 8</u>).

Parameter optimization was performed on a subset (training) of each dataset and the optimized parameters (Supplementary File 9) were compared to the default ones on the samples not used for training (validation subset). Figure 3 shows the optimization results at ROI level. In general, the optimization process improved sensitivity by slightly decreasing specificity. For panelcnDataset, sensitivity was increased by a higher margin driven by CODEX2, which increased its sensitivity by 58.6%. On the other hand, tools were not able to improve or showed small differences in the In-house MiSeq dataset (Supplementary File 11).

**Figure 3.**
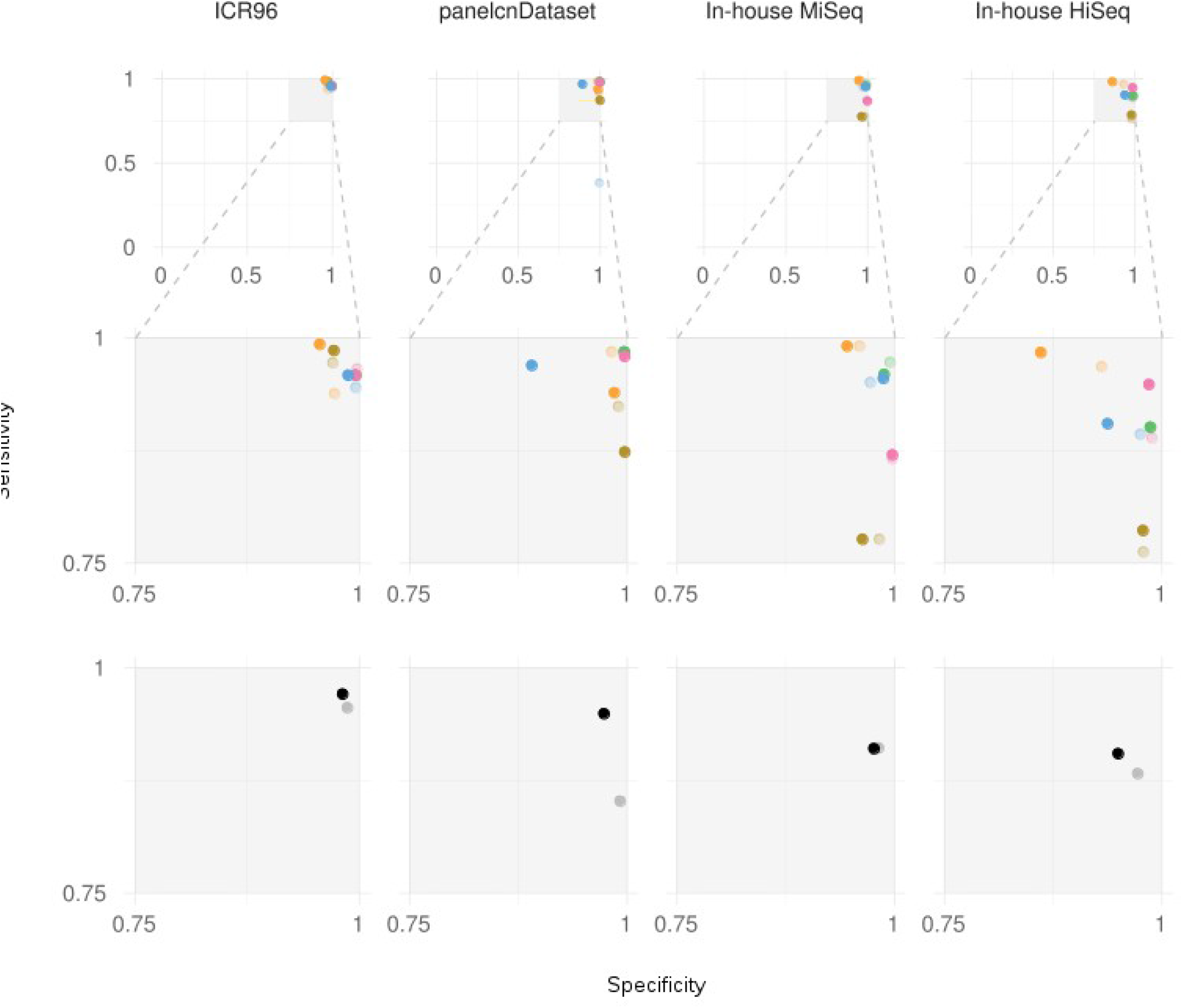
Optimization results at ROI level. Shows sensitivity and specificity on validation sets when executing tools with the optimized parameters in comparison to the default parameters.

### Benchmark in a diagnostics scenario

In a real diagnostic setting, all CNVs detected in genes of interest and all regions where the screening tool was not able to produce a result (no-call) should be confirmed by an orthogonal technique. To account for this, we evaluated the performance of all tools using the whole diagnostics strategy metric which takes the no-calls into account. This evaluation was performed in a modified version of the in-house datasets, the augmented in-house datasets (Figure 1), which contained all the samples from the original sequencing runs instead of a selection of them (see methods).

Figure 4 shows sensitivity and specificity on the augmented in-house datasets when executing tools with the optimized parameters compared to the default parameters. For the in-house MiSeq dataset, two tools detected all CNVs: panelcn.MOPS achieved it with both optimized and default parameters (80.2% and 67.1% specificity respectively), and DECoN only with the optimized parameters (90.8% specificity). CoNVaDING also detected all CNVs, but its high no-call rate led to very low specificity, 4.0%. For the in-house HiSeq dataset, only panelcn.MOPS detected all CNVs with an acceptable specificity: 81.7% and 83.3% with the default and optimized parameters, respectively. DECoN missed one CNV, being a mosaic sample, and its specificity remained high, 96.7% with the optimized parameters. On the other hand, CODEX2 and ExomeDepth obtained high sensitivity and specificity values for both datasets, but they did not report no-calls (Table 5 and Supplementary File 12).

**Figure 4.**
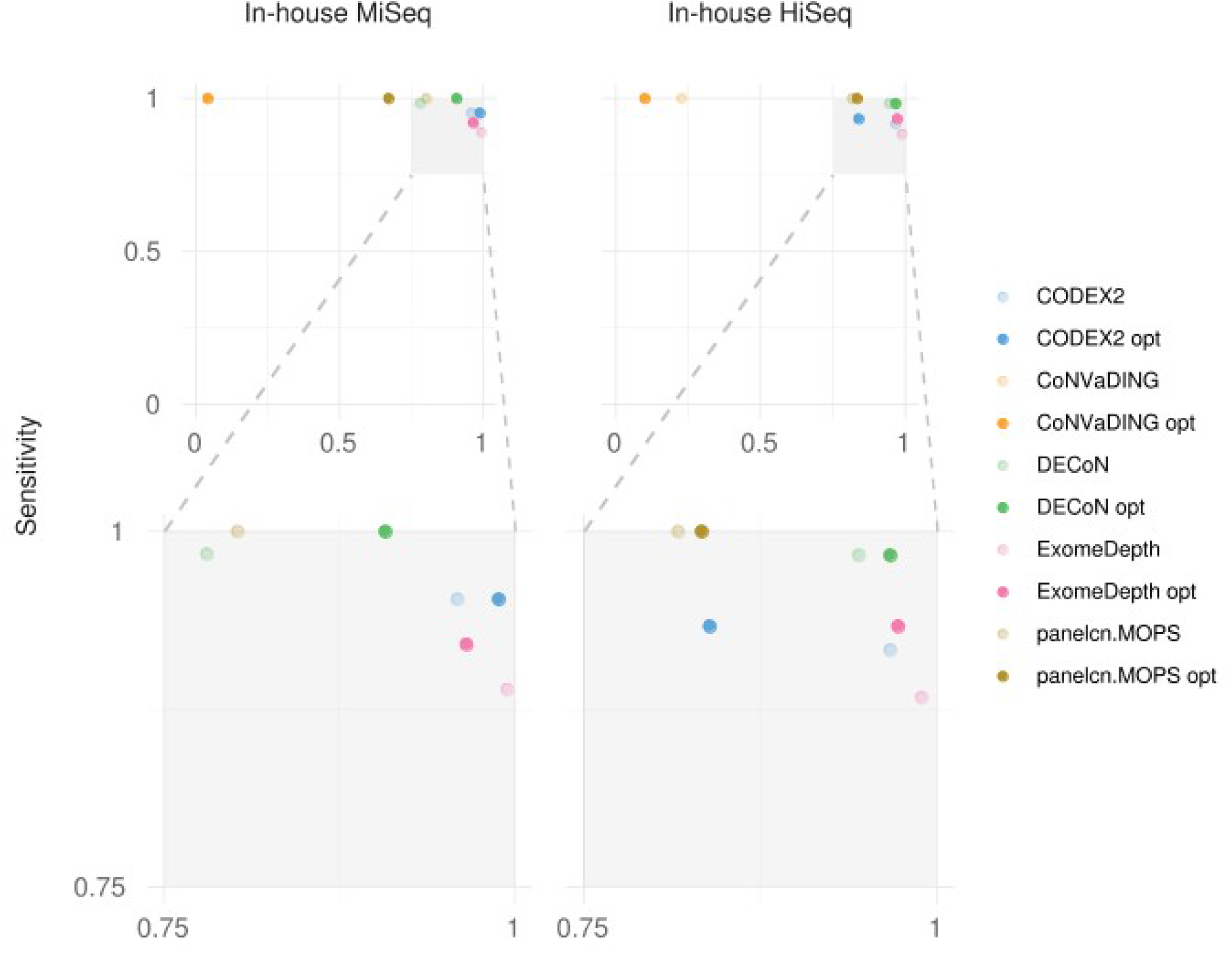
Benchmark results for the diagnostics scenario. Shows sensitivity and specificity on the augmented in-house datasets when executing tools with the optimized parameters in comparison to the default parameters.

## Discussion

CNVs are the genetic cause of multiple hereditary diseases (Zhang *et al*., 2009). To detect them, specific tools and techniques are required. In genetic diagnostics, this is mainly done using either MLPA and aCGH or using software tools to infer copy-number alterations from NGS data generated in the diagnostics process. MLPA and aCGH are the gold standard methods (Kerkhof *et al*., 2017), but both are time-consuming and expensive approaches that frequently lead laboratories to only use them in a subset of genes of interest. On the other hand, multiple tools for CNV calling from NGS data have been published (Zhao *et al*., 2013; Abel and Duncavage, 2013; Mason-Suares *et al*., 2016), but their performance on NGS gene panel data has not been properly evaluated in a genetic diagnostics context. This evaluation is specially critical when these tools are used as a screening step in a diagnostics strategy, since a non-optimal sensitivity would lead to a higher number of misdiagnosis.

Most CNV calling tools have not been developed to be used as a screening step in genetic diagnostics but as part of a research-oriented data analysis pipeline. Therefore, they were originally tuned and optimized for a certain sensitivity-specificity equilibrium. To be used as screening tools, we need to alter their default parameters to shift that equilibrium towards maximizing the sensitivity even at the expense of lowering their specificity. This parameter optimization must be performed in a dataset-specific way, since tools show performance differences between dataset due to dataset specificities coming from target regions composition, technical differences or sequencing characteristics.

In this work, we selected five tools that have shown promising results on panel data, and we measured their performance, with the default and sensitivity-optimized parameters, over four validated datasets from different sources: a total of 495 samples with 231 single and multi-exon CNVs. CNVbenchmarkeR, a framework for evaluating CNV calling tools performance, was developed to undertake this task. We also evaluated their performance in a genetic diagnostics-like scenario and showed that some of the tools are suitable to be used as screening methods before MLPA or aCGH confirmation.

### Benchmark with default parameters

The benchmark with default parameters showed that most tools are highly sensitive and specific, but the top performers depend on the specific dataset. Most tools performed the best when using data from panelcnDataset. DECoN, ExomeDepth and CoNVaDING reached almost 100% sensitivity and specificity. A possible reason for this is that this dataset contains the lowest number of single-exon CNVs (n=13), which are the most difficult type of CNVs to be detected. DECoN was the best performer for ICR96, a dataset published by the same authors, but other tools obtained similar results in that dataset. CoNVaDING was the most sensitive tool when analyzing our in-house datasets but showed the lowest PPV in all datasets with the exception of panelcnDataset. ExomeDepth showed the highest PPV in all datasets, making it one of the most balanced tools regarding sensitivity and specificity. Differences in tool performance depending on the dataset were also observed in previous works (Hong *et al*., 2016; Sadedin *et al*., 2018).

### Optimization

The different CNV calling tools included in this work were originally designed with different aims with respect to their preferred sensitivity and specificity equilibrium or the type of CNVs they expected to detect, and this is reflected in their default parameters and their performance in the initial benchmark. Our aim with this work was to evaluate these CNV callers as potential screening tools in a genetic diagnostics setting and for this reason, we required their maximum sensitivity.

The parameter optimization process allowed us to determine the dataset-specific parameter combination maximizing their sensitivity without an excessive specificity loss. The optimization had a different impact on different tools: while CODEX2 had shown a higher sensitivity in all four datasets the rest of the tools showed modest improvements. This is mainly due to the fact that sensitivity was already over 0.9 for most combinations and the number of false negatives to correctly call was small (between 4 and 8) in the per gene metric.

The final optimized parameters were dataset-specific, so we do not recommend using them directly on other datasets where the data has been obtained differently (different capture protocol or sequencing technologies, for example).

Based on our results, we would recommend optimizing the parameters for each specific dataset before adding any CNV calling tool to a genetic diagnostics pipeline to maximize its sensitivity and reduce the risk of misdiagnosis. To that end, we have developed an R framework, CNVbenchmarkeR (freely available at https://github.com/TranslationalBioinformaticsIGTP/CNVbenchmarkeR), that will help to perform the testing and optimization process in any new dataset.

### Diagnostics scenario

Two tools showed performance good enough to be implemented as screening methods in the diagnostics scenario evaluated in our two in-house datasets (Figure 4): DECoN and panelcn.MOPS. While panelcn.MOPS was able to detect all CNVs both with the default and the optimized parameters, DECoN reached almost perfect sensitivity and outperformed panelcn.MOPS specificity when using the optimized parameters. It only missed a mosaic CNV affecting two exons of the NF2 gene. CoNVaDING also reached 100% sensitivity, but the high number of no-call regions reduced its specificity to only 4%, which rendered it non-valid as a screening tool.

The parameter optimization process improved the sensitivity of most tools. For example, for the in-house MiSeq dataset, DECoN sensitivity increased from 0.98 to 1, and specificity increased from 0.78 to 0.91. This improvement highlights the importance of fine-tuning the tool parameters for each specific task, and shows that the optimization process performed in this work has been key for the evaluation of the different tools.

The high sensitivity reached by DECoN and panelcn.MOPS in different datasets, where they identified all known CNVs, shows that NGS data can be used as a CNV screening step in a genetic diagnostics setting. This screening step has the potential to improve the diagnostics routines. As an example, the high specificity reached by DECoN in the in-house MiSeq dataset with the optimized parameters means that 91% of genes with no CNV wouldn’t need to be specifically tested for CNVs when using DECoN as a screening step. The resources saved by the reduction in the number of required tests could be used to expand the number of genes analyzed, potentially increasing the final diagnostics yield.

## Conclusions

According to our analysis, among the five tested tools, DECoN and panelcn.MOPS provide the highest performance for CNV screening before orthogonal confirmation. Although panelcn.MOPS showed a slightly higher sensitivity in one of the datasets, DECoN showed a much higher specificity in the diagnostics scenario. Our results also showed that tools performance depends on the dataset. Therefore, it may be important to evaluate potential tools on an in-house dataset before implementing one as a screening method in the diagnostics routine.

## Acknowledgments

This study makes use of the ICR96 exon CNV validation series data generated by Professor Nazneen Rahman’s team at The Institute of Cancer Research, London as part of the TGMI. We are grateful to the Katharina Wimmer team at Division Human Genetics, Medical University Innsbruck for providing access to the dataset deposited at EGA and hosted by the EBI, under the accession number EGAS00001002481.

This work has been supported by: the Spanish Ministry of Science and Innovation, Carlos III Health Institute (ISCIII), Plan Estatal de I+D+I 2013–16, and co-financed by the FEDER program; the Government of Catalonia, the Spanish Association Against Cancer (AECC) and Fundació La Marató de TV3. Contract grant numbers: ISCIIIRETIC RD06/0020/1051, RD12/0036/008, PI11/1609, PI13/00285, PIE13/00022, PI14/00577, PI15/00854, PI16/00563, PI19/00553 and 2017SGR1282 and 2017SGR496.

We thank the participating patients and all the members of the Unit of Genetic Diagnostics of the Hereditary Cancer Program of the Catalan Institute of Oncology (ICO-IDIBELL) and the Genetics Diagnostics Unit of the Hereditary Cancer Group of the Germans Trias i Pujol Research Institute (IGTP). We also thank the IGTP HPC Core Facility and Iñaki Martínez de Ilarduya for his help.

