## Supplementary material for "Benchmark of tools for CNV detection from NGS panel data in a genetic diagnostics context": Supplemetal File 8

**Optimization algorithm**

A greedy approach was used to optimize the parameters of the algorithms to maximize sensitivity while limiting specificity loss or improving it when possible. For each dataset, we defined a training set with 50% of the samples to optimize algorithm parameters and a validation set with the other 50% to evaluate them.

The greedy algorithm behaves as follows. The default parameter, *D*, is used as reference. For each numeric parameter, 22 values between *D^0.25^* and *D^1.75^* are considered; only 9 values for CODEX2 were considered due to its high CPU requirement. For categorical parameters, all options are considered. Initially, the best solution is the execution with default parameters. Optimization starts from the first parameter: the algorithm is executed evaluating the whole values range for this parameter and keeping other parameters with default values. Solution with highest whole diagnostics strategy sensitivity is chosen if specificity decreases less than 30% in comparison to the previous best solution. If the whole diagnostics strategy sensitivity cannot be improved, then the highest per gene sensitivity is chosen if specificity decreases less than 25%. Finally, if per gene sensitivity cannot be improved, the highest per ROI sensitivity is chosen if specificity decreases less than 20%. In case of a tie at any level, the solution with the best specificity is considered. The best parameter value is therefore fixed. Optimization continues from a second parameter randomly chosen. The algorithm is executed evaluating the whole values range for this parameter and keeping other parameters with default values or fixed values. The process is repeated until the last parameter is reached.

To overcome the dependence of greedy algorithms on the order of parameter selection, the whole optimization process was repeated starting from each different parameter.

Below is shown the pseudocode describing the main steps for the optimization algorithm.

**Main steps of optimization algorithm**

best_solution = execution with default parameters

**for each** parameter p

D = default parameter value

**if** p is numerical:

values_range = 22 values from [D^0.25^, D^1.75^]

```
else if p is categorical:
	values_range = all categorical values
```

executions = ∅

**for each** v ∈ values_range:

v_previous = fixed values for already optimized parameters

v_next = default values for not optimized parameters

algorithm_parameters = v ∪ v_previous ∪ v_next

executions = executions ∪ execute_algorithm(algorith_parameters)

metrics = {whole strategy sensitivity, per gene, per ROI}

**for each** m ∈ metrics **and while** !success:

```
	local_best = highest_sensitivity(executions, m)
	coeff = 0.7 if (m = whole strategy sensitivity)
	coeff = 0.75 if (m = per gene)
	coeff = 0.8 if (m = per ROI)

if sensitivity(local_best, m) > sensitivity(best_solution, m)
	and specificity(local_best, m) > coeff * specificity(best_solution, m):
	best_solution = local_best
	success = true
	p is fixed with value from local_best
```
