## Supplementary material for "Benchmark of tools for CNV detection from NGS panel data in a genetic diagnostics context": Supplemetal File 1

**IBK141 exclusion from EGAD00001003335 dataset**

IBK141 sample is supposed to have an E54 NF1 deletion as described in the dataset EGAD00001003335. Using IGV for comparing our IBK141 bam file against other samples we see E54 coverage has a different shape:

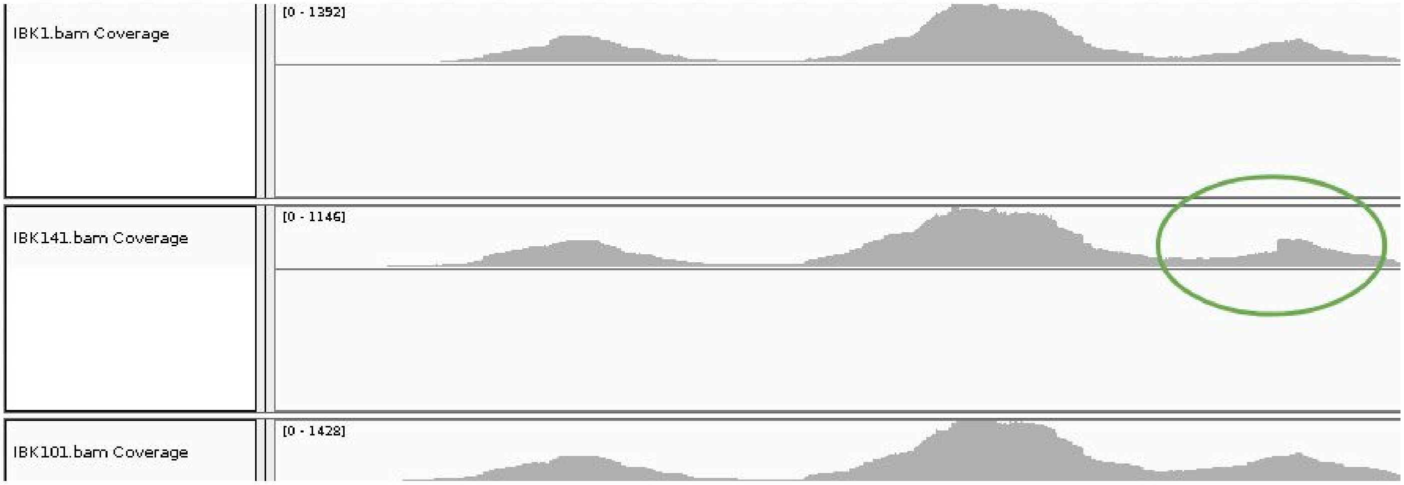


If we look at the mapped reads we observe that a good amount are soft clipped reads:

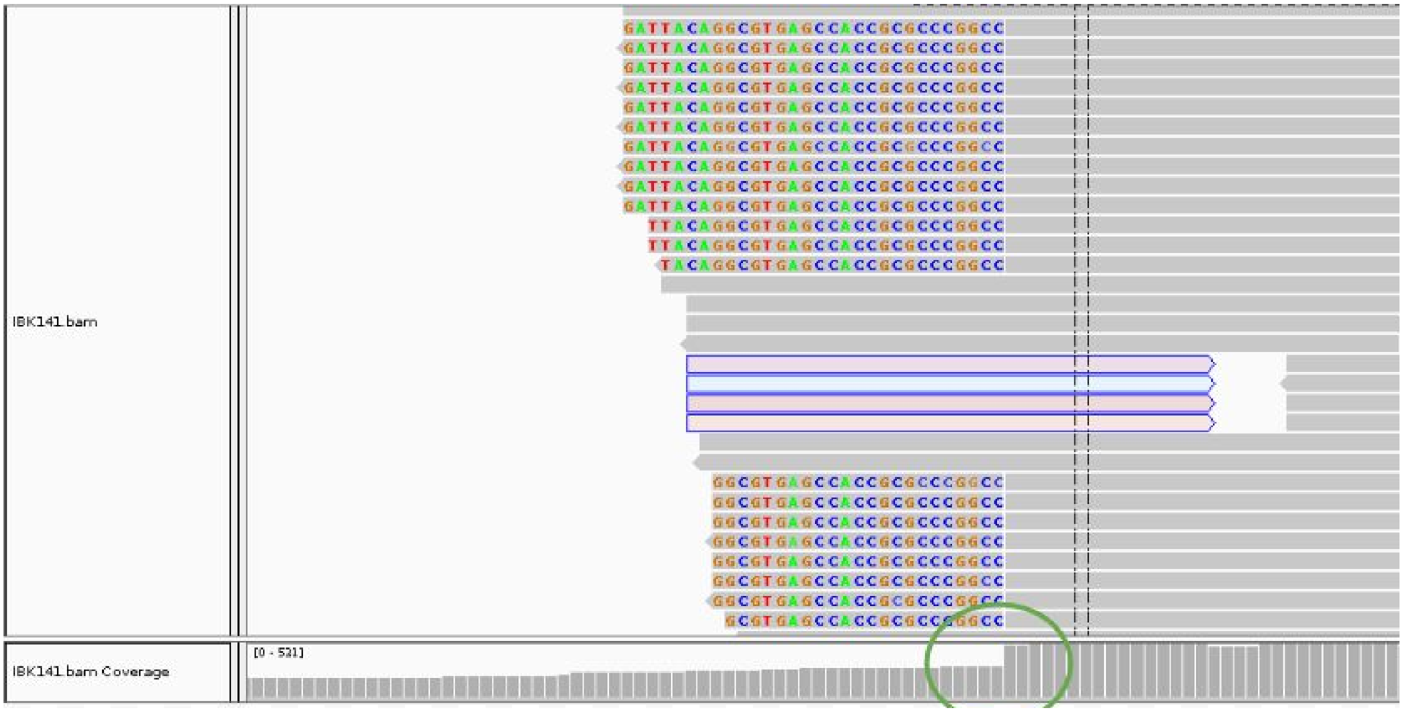


The soft clipped sequence in BLAST obtains 100/99% of identity with several hits for primates which may indicate that this sequence correspond to an ALU.

Additionally, if we look at the start of the **NF1 MLPA probe**, we see that the clipped regions end just in the middle of the reference sequence recognized by the MLPA probe (AGTTT**GATCAAC**):

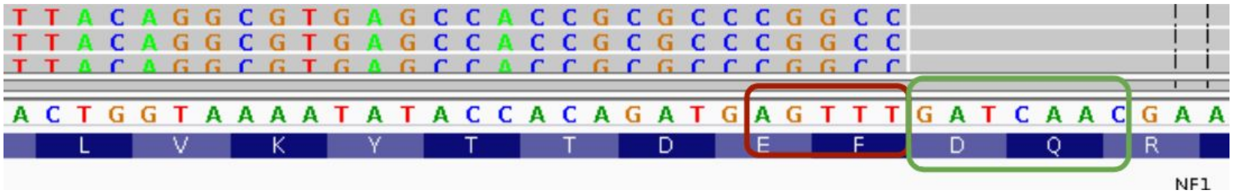


Accordingly, our hypothesis is that there wasn't any exon deletion: an ALU was inserted just in the middle of the MLPA probe start sequence, so the MLPA can fail when recognizing the exon as deleted due to the coincidence between the probe start and the point where the ALU was inserted. For this reason, IBK141 sample was removed from the the EGAD00001003335 dataset.
